# Dance displays in gibbons: Biological and linguistic perspectives on structured, intentional and rhythmic body movement

**DOI:** 10.1101/2024.08.29.610299

**Authors:** C. Coye, K.R. Caspar, P. Patel-Grosz

## Abstract

Female crested gibbons perform conspicuous sequences of twitching movements involving the rump and extremities. However, these dances have attracted little scientific attention and their structure and meaning remain largely obscure. Here we analyse close-range video recordings of captive crested gibbons, extracting descriptions of dance in four *Nomascus* species (*N. annamensis, N. gabriellae, N. leucogenys,* and *N. siki*). Additionally, we report results from a survey amongst relevant professionals clarifying behavioural contexts of dance in captive and wild crested gibbons. Our results demonstrate that dances in *Nomascus* represent a common and intentional form of visual communication restricted to sexually mature females. While primarily used as a proceptive signal to solicit copulation, dances occur in a wide range of contexts related to arousal and/or frustration in captivity. A linguistically informed view of this sequential behaviour demonstrates that dances follow a grouping organisation and isochronous rhythm – patterns not described for visual displays in other non-human primates. We argue that applying the concept of dance to gibbons allows us to expand our understanding of the communicative behaviours of non-human apes and develop hypotheses on the rules and regularities characterizing this behaviour. We propose that gibbons dances likely evolved from less elaborate rhythmic proceptive signals, similar to those found in siamangs. Although dance displays in humans and crested gibbons share a number of key characteristics, they cannot be assumed to be homologous. Nevertheless, gibbon dances represent a model behaviour whose investigation could be extended to the study of complex gestural signals in hominoid primates.

## 1. INTRODUCTION

### 1.1 BACKGROUND AND AIM OF THIS STUDY

The concept of *dance* is primarily discussed in the context of human communication, where dancing is defined as intentional, rhythmic and non-mechanically effective body movements (see e.g. Hanna 1979, 2017). However, this definition can also be applied to certain social behaviours in other animals, including non-human primates, whose (potential) dance displays have only received limited attention so far (Francis 1991; Fan et al. 2016 for gibbon dances; Bertolo et al. 2021 for chimpanzee rhythmic displays). Indeed, it overlaps with that of a communicative *gesture* in non-human primates, typically defined as a discrete, intentional movement of a body part, which is potentially detectable by an audience and non-mechanically effective (Genty et al. 2009). Notably, dance is broader, in that it includes non-discrete movements and ‘global’ movements of the entire body, and definitions of gesture generally lack the rhythmic component central to common definitions of dance (as in the anthropological work of Hanna 1979, 2017). Criteria used to assess intentionality in animal gestures and dance (inherent to both definitions) vary between studies but usually include sensitivity to the presence or attentional state of an audience, persistence and/or elaboration until the emitter’s goal is reached (Cartmill and Byrne 2010; Genty et al. 2009).

Here we focus on an understudied form of visual communication that is characteristic to adult female small apes of the genus *Nomascus*, commonly referred to as crested gibbons. Our approach relies on the joint efforts of linguists and biologists, aiming to characterise complex animal communication signals via means of both biological and linguistics tools (i.e. see Berthet et al. 2023). The visual display studied here takes the shape of rhythmic intentional movements of the entire body, thus qualifying as dance, and as different from gesture.

The central insight from our study is that gibbon dance displays systematically qualify as an intentional and rhythmic communicative body movement that follows a non-random structure, which can be captured in terms of grouping structure as previously proposed for human dance. This novel description fills a gap in our understanding of the communicative behaviours of non-human apes. *Dance* originates as a concept specific to human culture and interaction, which should only be applied to other species when a comparison is of benefit for scientific discovery. We show that this is the case, as an application of tools that have previously been developed for human dance (such as grouping analysis) allow us to develop hypotheses on the rules and regularities that characterize the gibbons’ dance behaviour.

### 1.2 DANCE DISPLAYS IN GIBBONS

Gibbons are exclusively arboreal apes (superfamily Hominoidea) that live in small territorial social units, typically structured around a single monogamous breeding pair (Malone and Fuentes 2009). With regards to social communication, investigations have largely focused on their vocal behaviour, which include highly elaborate song bouts (Clarke et al. 2006; Geissmann 2002). *Nomascus* represents one of four extant gibbon genera (Mootnick and Fan, 2011). Taxonomists distinguish seven *Nomascus* species, which occur in the tropical and temperate forests of Southern China, Eastern Indochina and the island of Hainan (Mootnick and Fan 2011; Roos 2016). All of these species are sexually dichromatic, with dark pelage colour in males and light colouration in females, and exhibit striking ontogenetic colour change (Mootnick and Fan 2011). At the behavioural level, *Nomascus* differ from other small apes in the frequent occurrence of polygynous rather than monogamous groups in wild populations (Delacour 1933; Guan et al. 2018; Li et al. 2022). However, polygynous habits have so far mostly been reported from northern *Nomascus* species (*N. concolor, N. hainanus, N. nasutus*), while the more limited number of studies on the socioecology of the southern species group (*N. annamensis, N. gabriellae, N. leucogenys, N. siki*) suggest polygyny to be rather exceptional (Barca et al. 2016; Hu et al., 1989; Kenyon et al. 2011). The diversity and usage of facial expressions in crested gibbons is similar to that of other hylobatids (Florkiewicz et al. 2018; Scheider et al. 2016); their gestural repertoire has never been comparatively assessed in a rigorous manner (but see De Vries, 2004).

Dances in crested gibbons were first noted anecdotally in captive individuals (Maxwell, 1984). Lukas et al. (2002) were the first to systematically monitor the occurrence of such displays (therein described as “bobbing”), remarking that their frequency increased during oestrus in the single mated Northern white-cheeked gibbon (*N. leucogenys*) female that they studied. Subsequent observations documented the occurrence of dances in wild crested gibbons and support the assumption that they function primarily as proceptive signals to solicit copulation and are only displayed by adult females (Fan et al. 2016; Li et al. 2022; Zhou et al. 2008). During a dance, the gibbons do not vocalise (Fan et al. 2016).

To date, the most comprehensive analysis of this behaviour has been provided by Fan et al. (2016), who report the occurrence of dances in four wild adult females of the Eastern black crested gibbon (*N. nasutus*). They characterise the female display in terms of “a rhythmic moving of her body (arms, legs, trunk, and head) […] while maintaining other body parts motionless […] similar to a human ‘Robot Dance’” (Fan et al. 2016). The authors offer observations of the contexts in which dances occur and also provide descriptions of the structure of two selected dancing bouts. Finally, they noted the subsequent behaviour of associated males, who responded positively (approach, grooming or copulation) to 46.2% of dancing bouts (112 of 242 instances). Fan et al. (2016) hypothesise that dancing serves several functions: besides soliciting copulation, it may also strengthen inter-sexual social bonds and could represent a form of non-aggressive intrasexual competition between females within polygynous groups. However, although *Nomascus* dances are frequently observed in both wild and captive settings (Lukas et al. 2002; De Vries, 2004; Burns and Judge 2015; Fan et al. 2016; pers. obs.), neither rigorous tests of these hypotheses nor detailed analyses on the phylogeny, structure, and variability of dances have been conducted.

In this study, we analysed close-range video recordings of captive crested gibbons to provide novel descriptions of dance in females from four *Nomascus* species (*Nomascus annamensis, N. gabriellae, N. leucogenys* and *N. siki*). We focus on three aspects that have previously not been explored: grouping structure of movements, intentionality, and rhythmicity. These aspects of dance are hard to study in the wild, because the canopy habitat of gibbons can easily obstruct an observer’s view. In addition, we report the results from a survey among professionals on the occurrence and context of dance displays in captive as well as wild crested gibbons.

## 2. MATERIALS AND METHODS

All statistics were performed in R (R Core Team, 2023).

### 2.1 DATA COLLECTION AND DEFINITION OF GIBBON DANCE

In total, we compiled 37 videos capturing behavioural sequences including a dance event, defined as an abruptly commencing temporary stiffening of the body accompanied by rhythmic, often repetitive twitching body movements (e.g. movement of the rump and/or the limbs and head). To illustrate the definition of dance event, consider the homogeneous dance in Supplementary File 1 (starting at time stamp 02:29); the stiffening of the body commences with the beginning of the video (02:33), and ends after 19 seconds (02:52), when the gibbon turns her head and relaxes the body; the twitching body movement in this video occurs in approximately one-second intervals, yielding a total of 19 visible twitches. Movements that did not fit this definition but appeared within a dance sequence (self-scratching; locomotion) were excluded from the analyses. In the vast majority of dance sequences, the dancing female’s back was turned towards the recipient, but apart from this general trend, dance structure was highly variable.

Out of the 37 initial videos, 11 were discarded from all subsequent analyses due to external disturbances (e.g. a human touching the animal). The 26 remaining videos were used for the intentionality analysis. Seven of those videos were further discarded from the grouping analysis due to limited visibility preventing a fine-grained analysis of movement patterns and five were removed from the rhythm analyses because they did not meet the respective criteria (see below). The videos were opportunistically recorded from 7 captive *Nomascus* females housed in European and Australian zoos (Table 1), as well as in the Endangered Primate Rescue Centre (EPRC Vietnam). Information about the identity and origin of each subject in the video analysis are summarised in Table 1. Video material was either derived from the personal archives of the authors or was solicited from researchers and zoo staff via the research survey accompanying this study. Videos from Zoo Duisburg were originally recorded for a previous cognitive study on gibbons (Winking, 2016), capturing the behaviour by chance. Apart from the footage from Duisburg, each video captured just one dance event.

**Table 1:** Information on subjects and dance sequences considered for analyses on intentionality and grouping (EPRC: Endangered Primate Rescue Center, Cúc Phương, Vietnam). The column “total number of dances” equals the number of dances considered for the intentionality analysis.

| Species | Individual | Location | Age in years at recording | total # of dances | # of dances in grouping analysis | # of dances in rhythm analysis |
| --- | --- | --- | --- | --- | --- | --- |
| *Nomascus annamensis* | Hu (TJD21-00751) | EPRC | ~ 9 (wildborn) | 7 | 7 | 7 |
| *Nomascus gabriellae* | Ina (TJD19-00500) | EPRC | 22 | 9 | 8 | 8 |
|  | Lucki (MIG12-29370588) | Zoo Duisburg | 7 | 12 | 0 | 5 |
|  | Polly (TJD19-00461) | EPRC | ~ 14 (wildborn) | 1 | 0 | 1 |
|  | Jermei (MIG12-29965298) | Perth Zoo | 21 | 1 | 1 | 1 |
| *Nomascus siki* | Doremon (TJD19-00497) | EPRC | ~ 7 (wildborn) | 2 | 2 | 2 |
|  | Hope (TJD19-00491 ) | EPRC | 11 | 1 | 1 | 1 |

Observers unfamiliar with the behaviour of crested gibbons might interpret dances as a kind of locomotor stereotypy, potentially indicative of poor welfare. However, behavioural stereotypies in gibbons are well documented and include excessive scratching, body rocking, repetitive brachiation, masturbation, and self-inflicted harm (Mootnick & Baker, 1994; Cheyne, 2006; Cooke & Schillaci, 2007; Hosey & Skyner, 2007). All of those behaviors are highly distinct from dances and, in contrast to them, are absent in wild populations. In captivity, stereotypies have been reported from both sexes, while dances appear to be restricted to sexually mature females (see below).

In many of the analysed videos (e.g., see Supplementary File 1) the gibbons move their body along the wire mesh of a fence while dancing. We want to emphasize that this fence rubbing is a side effect of the gibbons targeting their dance at a human observer on the other side of the barrier and moving close to them. The dance itself does not represent a form of purposeful self-scratching. Crested gibbons use their hands and less frequently their feet to groom through their fur instead of rubbing their limbs or backs against a substrate (e.g., Mootnick et al. 2012). Moreover, fence-rubbing during dancing would not be mechanically effective, especially during homogeneous displays (see below).

### 2.2 ASSESSMENT OF INTENTIONALITY

We used standard criteria from great ape research to assess intentionality (Cartmill and Byrne 2010; Genty et al. 2009): sensitivity to the attentional state of the audience (assessed via measures of audience checking and attention-getting behaviours), persistence (i.e., pursuit of the behaviour after audience checking) and elaboration (i.e. inclusion of novel behaviours to the dance). The presence or absence of each of these behaviours was assessed by an experienced coder (CC) and scored as a binary variable for each dance bout. All cases in which the applicability of these concepts was deemed to be even slightly ambiguous have been coded as negative, thus leading to a highly conservative assessment. An audience (conspecific and/or heterospecific) was present and able to see the dancing female in each of the videos analysed. A potential caveat is that more than half of the dances were recorded by an experimenter holding a camera in close proximity to the gibbon, thus making the presence of an attentive audience (i.e. an individual oriented towards the female) a pre-requisite for the acquisition of the footage. This is one of the reasons why a more fine-tuned criterion to assess audience sensitivity (i.e. audience checking) was adopted.

### 2.3 ASSESSMENT OF STRUCTURAL GROUPING

*Nomascus* dance displays involve rhythmic body movements, which sets them apart from the gestures of non-human great apes, and makes them more similar to human dance. To lay the groundwork for a better understanding of gibbon dance displays, we capitalize on the similarity between gibbon dance and human dance, and ask whether gibbon dance follows structural regularities of the type that has been proposed for human dance (Charnavel, 2019), amounting to a rudimentary ‘dance grammar’. In the analysed videos, gibbon dance movements can often be described as left-right, up-down, or diagonal movements (see Table 2 for details), and left-right movement occurs both in a sitting posture and in a standing posture. Current approaches to the structural organization of human dance build on segmentation/grouping (building on Wertheimer 1938, Lerdahl & Jackendoff, 1983, amongst others), where similar behaviours are grouped together, and behaviour changes give rise to a group boundary. Based on that, we qualitatively analysed the structure of the dancing bouts in terms of *grouping*. Analogous to research on human dance (Charnavel, 2019), groups were defined as homogeneous, continuous behavioural sequences which constitute unitary blocks within the dance bout. Grouping, when attested, gives rise to a (rudimentary) syntax, i.e., a system of possible movement sequences that is plausibly governed by rules that generate well-formed and ill-formed sequences (compare Berthet et al. 2023).

**Table 2:** Behavioural variables included in the grouping analysis and their definition.

| Behavioural category | Variable | Definition |
| --- | --- | --- |
| Global body movement | Up-down | Repeated vertical movement of the body |
|  | Left-right | Repeated lateral shifting of the body |
|  | Diagonal | Repeated diagonal movements of the body (i.e. the direction cannot be coded as purely vertical or horizontal) |
|  | Front-back | Repeated body movement from front to back, with no vertical or lateral component |
|  | Body shake | Stereotyped shaking of the entire body (similar to a gibbon's movement to rid its coat of water – Baldwin & Teleki, 1976) |
| Movement of discrete body parts (limbs or head) | Head movement | The individual turns its head to either side |
|  | Arm extension | The individual extends an arm away from the body * |

\* This behavioural item does not include begging, which we define as “sustained extension of an arm in the direction of a human observer, with clear eye contact”

We state three theoretical hypotheses for grouping in gibbon dances and illustrate the predictions for these three movement types (where LR stands for ‘left-right’, UD for ‘up-down’ and DIA for ‘diagonal’):

**Null hypothesis (no grouping):** movement sequences are random; each dance movement (twitch) is independent from the preceding dance movement. **Prediction:** no discernible pattern; e.g., we would expect to find sequences such as LR-UD-DIA-LR-DIA-UD.

**Alternative hypothesis 1 (one-level grouping):** dances are segmented into groups of similar dance movements, exhibiting a one-level grouping structure. **Prediction:** discernible patterns where group boundaries can be established on the basis of changes in movement parameters; e.g., we would expect to find sequences such as [LR-LR-LR]-[UD-UD-UD].

**Alternative hypothesis 2 (two-level grouping):** dances are segmented into groups of similar dance movements, exhibiting a two-level grouping structure, where some groups can be grouped together further based on a higher-level similarity. **Prediction:** discernible patterns where group boundaries can be established on the basis of changes in movement parameters, but some changes have a more intense effect than others; e.g., we would expect to find sequences such as [[LR_standing_-LR_standing_-LR_standing_]-[LR_sitting_-LR_sitting_-LR_sitting_]]-[[UD-UD-UD]], where change in direction would have a more intense effect than change in posture.

Groupings of movement patterns in the gibbons’ displays were assessed by a trained observer (CC). In addition to the subject’s posture (i.e. sitting, standing, hanging), selected behavioural variables were coded systematically (Table 2) and used to define groups of movements and group boundaries within a display bout.

Finally, we computed inter-observer comparisons using data from a second coder (KRC, who recoded ∼20% of the dataset) using Cohen’s kappa to assess the match for scores of postures and directions of movement as well as intra-class correlations (model: two-way; type; agreement) for the length of the groups (measured in seconds). These tests confirmed a good match between observers (Direction: κ = 0.79; Posture: κ = 0.75, Groups duration: ICC = 0.787; *F*-test *p* < 0.001).

### 2.4 RHYTHM

The body twitches of a dancing gibbon appear rhythmic, but their temporal structure had so far not been quantified. To accomplish this, we adopted the methodology of De Gregorio et al. (2023), who recently studied rhythm in the songs of *Nomascus* gibbons, and applied it to the temporal occurrence of twitches within dance sequences. Videos were loaded into the software BORIS (Friard & Gamba, 2016) and viewed at 10% natural speed to code onsets of a twitch movement. Based on this, we determined the inter-onset intervals (IOI) as well as rhythmic ratios (*r*) of consecutive IOIs within each dance. An IOI was defined as the time duration between the onsets of two consecutive twitches within a dance sequence. We calculated *r* with the formula IOI*_n_*/(IOI*_n_+*IOI*_n_*_+1_) where *n* is the position of the respective IOI in a dance sequence (De Gregorio et al., 2023). Only dances during which the respective gibbon was fully visible throughout the dance duration and which included at least 10 twitches were considered for rhythm analysis. As in De Gregorio et al. (2023), IOI ≥ 5 s were ignored, as were those that included body shakes (see Tab. 2) or brachiation bouts. Dances interrupted by one of these events were counted as separate sequences of movements.

We tested for interindividual differences in IOI and *r* using the Kruskal-Wallis test. Of special interest to us was the question whether gibbon dances, similar to the songs of at least some species (De Gregorio et al., 2023; Ma et al., 2024), follow an isochronous rhythm. Isochrony is given when consecutive IOI are of the same length, which is indicated by *r* = 0.5 (De Gregorio et al., 2023). To test for the presence of isochronous movement patterns, we visually inspected the data and ran Bonferroni-corrected two-sided one-sample Wilcoxon tests assuming a hypothetical median m_0_ = 0.5. Non-significant test outcomes would indicate the presence of isochrony, while significant deviations from the hypothetical median would suggest that our empirically determined rhythms would follow a non-isochronous pattern. We first tested the *r* distribution of each dance independently. Later, we also did so with a pooled *r* dataset comprising all quantified sequences after ensuring that there were no significant interindividual differences in this variable (Kruskal-Wallis test: χ²= 0.74776, *p* = 0.9934). We calculated single score intra-class correlations (model: two-way; type; agreement) for the twitch onsets of two dances (*n* = 90), which were quantified by two raters and yielded an excellent agreement (ICC = 0.943; *F*-test *p* = 0.0016). The rhythm dataset is included in Supplementary File 2.

### 2.5 RESEARCH SURVEY

We conducted a self-administered online survey (via Google Forms) which was disseminated by email to relevant professionals with expertise in crested gibbon behaviour (field primatologists and zoo staff, including curators, veterinarians and primate keepers). The distribution of the survey was approved and facilitated by the Gibbon Taxon advisory group (TAG) of the European Association of Zoos and Aquaria (EAZA) and the Gibbon Species Survival Plan® (SSP) steering committee of the Association of Zoos and Aquariums (AZA) after a careful evaluation of the project. The survey was made available between November 2022 and June 2023. It included 12 questions (Suppl. File 3), pertaining to the occurrence of dance, the sex, age and contraceptive status (i.e., with or without hormonal contraception) of the dancing individuals, the behavioural context of dances, and the professional background and experience of the respondent. The survey contained an exemplary video of a complex dance (Suppl. File 1) showcasing what type of behaviour the study dealt with. We received 29 responses to the survey, all of which were complete in the sense that all applicable questions were answered by the respondents.

## 3. RESULTS

### 3.1 INTENTIONALITY

We found dance in *Nomascus* to comply with criteria of intentional communication. In 59% of dances (i.e. 20 instances), a clear audience-checking behaviour during the dance could be identified (in 13 cases i.e. 38%, that behaviour was repeated several times during the dance bout). We identified 4 instances in which the female displayed attention-getting behaviours (3 instances in which the female repositioned herself in space to be in front of the receiver after she moved, and one hand slapping behaviour). Persistence after audience checking occurred in 53% of the dances (i.e. 18 dances). Finally, we found one case which could reflect elaboration: the females extended her arm in a begging gesture immediately after they stopped dancing. This behaviour needs to be interpreted carefully, as the contextual information available does not allow us to confirm that the female’s initial goal was to obtain food from the human towards whom the dance was directed.

### 3.2 GROUPING

The 19 dances analysed for grouping structure had a mean duration of 20 s (+/- 23, range: 6 – 126 s). Multiple groups were identified in 13 out of 19 videos analysed. Two main behaviour changes in gibbon dance sequences were observed that plausibly give rise to group boundaries (*Grouping Preference Rules, GPRs*), namely *change of contact point with the substrate/weight shift* (Charnavel’s GPR3) and *change of direction* (GPR4), as defined in Charnavel (2019:4). To elaborate, in GPR3, the contact point with the substrate changes (e.g. between sitting and standing) while the direction stays constant (e.g., left-to-right), a change in human dance sequences that would give rise to the perception of group boundaries. Similarly, a change from left-right movement to repeated rhythmic up-down movement would correspond to *change of direction* (GPR4), as the movements differ in the direction of their respective path. It is unclear whether these two rules are weighted differently when both create grouping boundaries in a given dance sequence. The two movement parameters are particularly suitable for a grouping analysis of gibbon dance, since they cover the two main categories of dance features (Charnavel, 2019:5): aspects of the posture while dancing (includes GPR3), and aspects of the dance movement (includes GPR4).

We recognise a spectrum of complexity in gibbon dances (Suppl. File 1). Within it, we identify three main types of grouping structures: homogeneous dances (*n* = 6), where the dance consisted of a single group; simple dances (*n* = 9), where we identified groups within the dance but without discernible further structure; and complex dances (*n* = 4), i.e., in which some group boundaries appear more *intense* than others, indicating a two-level grouping where smaller (lower-level) groups are nested within larger (higher-level) groups. Homogeneous dances (Fig. 1a) were expressed as rhythmic twitches or bobbing movements of the whole body and occurred without the sequential changes in posture seen in simple and complex dances. Simple dances (Fig. 1b) contained 4.4 groups on average (+/- 1.3), while complex dances (Fig. 1c) counted 7.25 groups on average (+/-1.7), which potentially qualify as lower-level groups in a structure with 4.5 higher-level groups (+/- 2.5)).

**Fig. 1:**
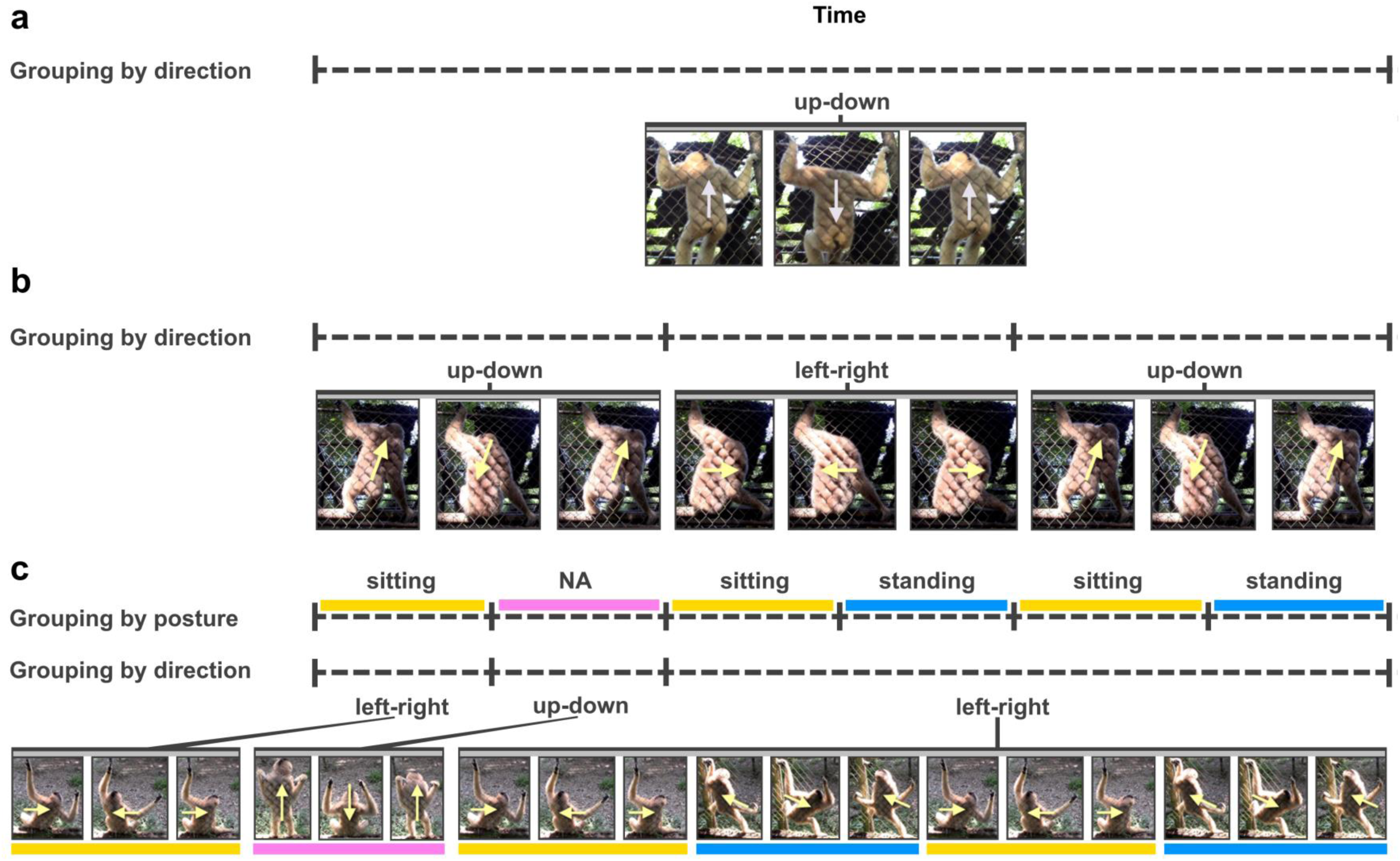
Schematic representation of movements in *Nomascus* dances. **a**: homogeneous dances. **b**: simple dances. **c**: complex dances. In the latter, postures are marked with colours. Note the alternation of non-nested left-right and up-down movements the first half, followed by left-right movements nesting in alternating sitting and standing postures

Repetition of dance movements occurs across all dance displays, and dance movements (twitches) do not exhibit the random choice of dance movement predicted by the null hypothesis (no grouping) in Section 2.3. Our observations thus militate against this null hypothesis. Alternative hypothesis 1 (one-level grouping) is sufficient to account for the majority of dances (*n* = 15), namely the homogenous dances and the simple dances. Complex dances are consistent with both alternative hypothesis 1 (one-level grouping) and alternative hypothesis 2 (two-level grouping), raising the possibility that two-level grouping occurs in gibbon dance. Fig. 1c illustrates two grouping analyses: grouping by posture gives rise to 6 groups; grouping by direction of movement gives rise to 3 groups. Since posture change clearly gives rise to group boundaries, a one-level grouping analysis (alternative hypothesis 1) assumes that only posture gives rise to a grouping structure (see Fig. 2a); however, the relationship between posture and direction seems non-random: from 00:27 until 01:10 in the recording, the gibbon consistently performs a left-right movement, while changing posture three times. An analysis with two-level grouping (alternative hypothesis 2) capitalizes on the observation that direction change seems to be the structuring principle of the entire dance, which means that posture change gives rise to lower-level grouping, while direction change gives rise to higher-level grouping (see Fig. 2b). The opposite is not feasible in Fig. 1c, as there are no instances where the same posture is maintained across a direction change. Since existing recordings of dance displays are relatively short, more data collection is needed to reveal whether there is a higher-level organization that follows systematic principles (alternative hypothesis 2; Fig. 2a), or whether only the lower-level organization exists (alternative hypothesis 1; Fig. 2a).

**Fig. 2:**
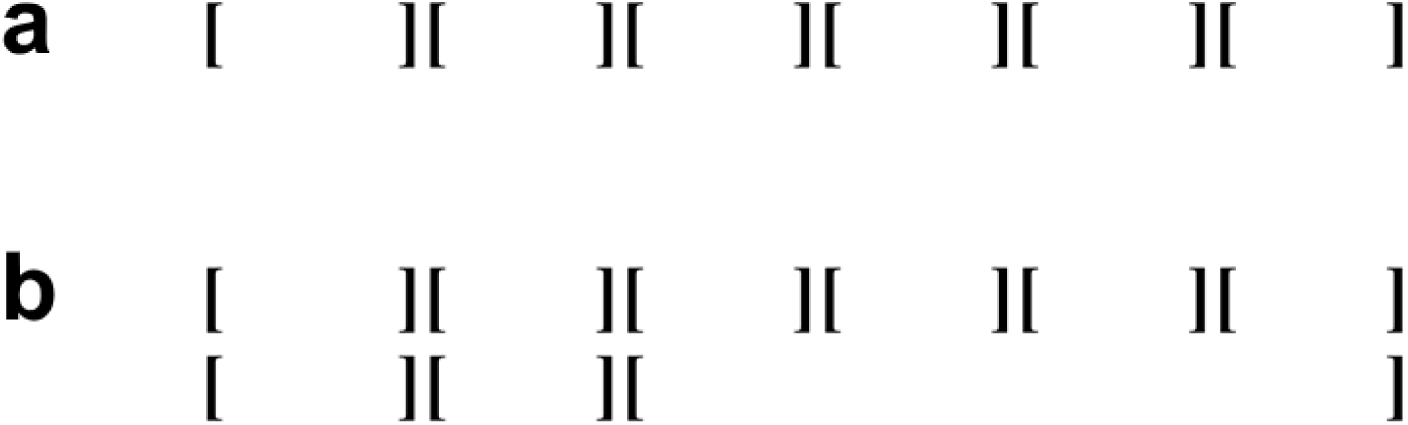
Two possible grouping analyses of a complex dance movement as illustrated in Fig. 1c. **a**: One-level grouping based on posture. **b**: Two-level grouping based on posture at the lower level and on direction at the higher level.

Intensity (of the effect of a behaviour change) is the guiding principle for positing higher-level group boundaries in line with alternative hypothesis 2 (Charnavel, 2019:17). For the complex dances (*n* = 4), we observed variation in whether posture change or direction change appears to have the more intense effect. In one complex dance, GPR3 (posture; change of contact point with the substrate) had a more intense effect; GPR4 (direction of (jerky) movement) had a more intense effect in two complex dances (including Fig. 1c); and in one case, each of GPR3 and GPR4 had a more intense effect at different parts of the dance. Interestingly, the choice of whether GPR3 or GPR4 had a more intense effect (thus potentially setting a higher-level group boundary) could change between dances, even within an individual. For instance, in one of Ina’s (*Nomascus gabriellae*, Endangered Primate Rescue Centre, see Table 1) dance sequences, posture had a more intense effect than direction of movement, and in other dance sequences it was the contrary.

A full analysis of the complex dance that underlies Fig. 1c is included in Supplementary File 4 and corresponds to the complex dance video in Supplementary File 1.

### 3.3 RHYTHM

We analysed 27 dances for rhythm patterns, quantifying a total of 1113 inter-onset intervals (IOI) between twitch movements within dance sequences. The length of IOI varied between 0.167 s and 3.203 s with a mean of 0.903 s (SD: 0.341). IOI length differed significantly between individuals (Kruskal-Wallis test: χ²= 448.92, *p* < 0.0001; Fig. 3a) but the rhythm of consecutive movements, quantified by the rhythmic ratio *r* (*n* = 1086), did not (Kruskal-Wallis test: χ²= 0.748, *p* = 0.993). A density plot of the pooled *r* dataset (Fig. 3b) highlights a unimodal distribution of values clustering conspicuously around 0.5 (mean: 0.5004). *r* = 0.5 indicates that consecutive IOI tend to be of equal length, in other words, the dances follow a predominantly isochronous pattern, regardless of their IOI length. In line with this, the distribution of *r* did not differ significantly from a hypothetical median of 0.5 in any of the individual dances (*p* > 0.480), nor in the pooled dataset (*p* = 0.658), suggesting isochronous dance rhythms.

**Fig. 3:**
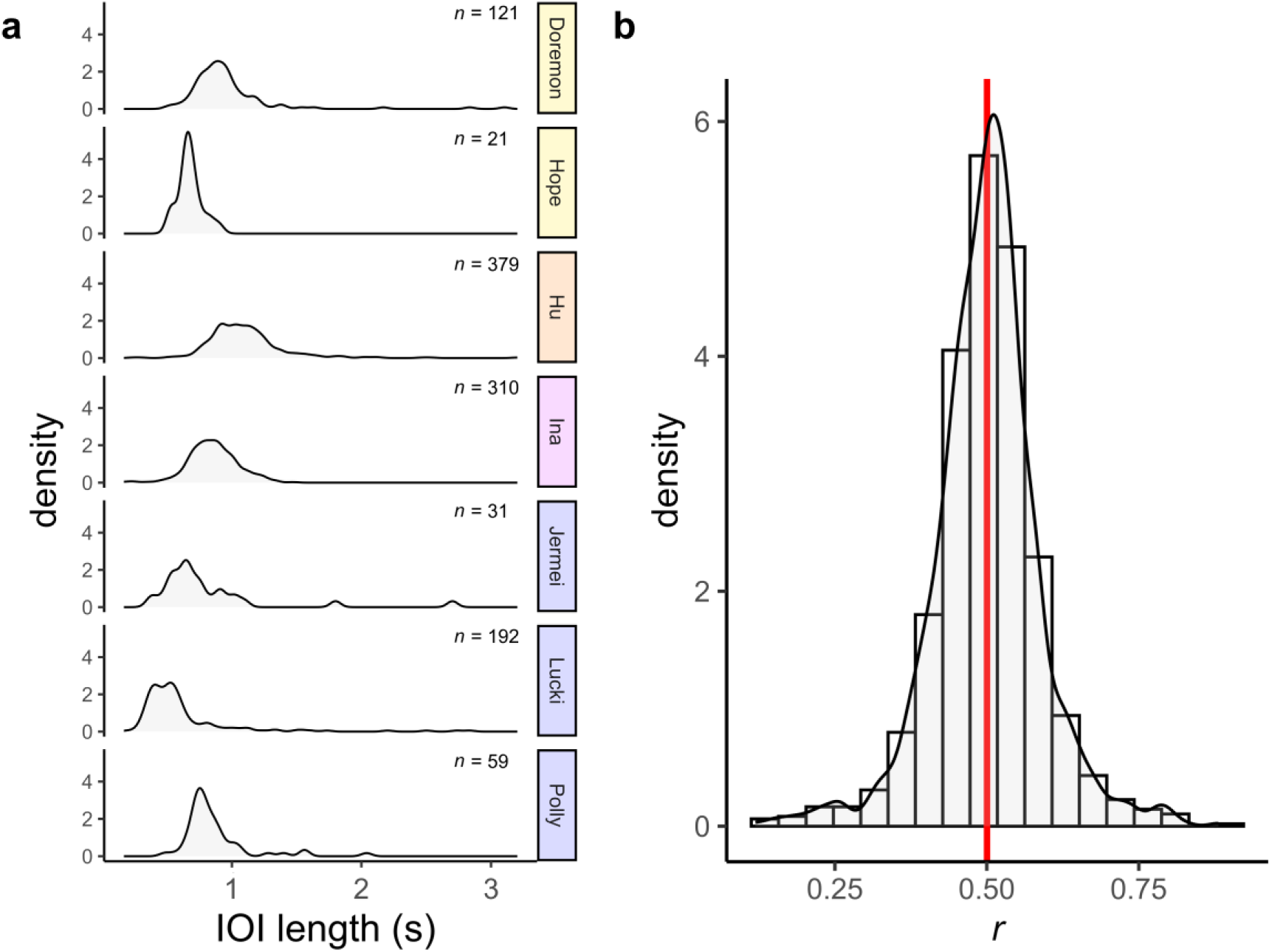
Rhythmic variables of dance in crested gibbons (*Nomascus* spp.). **a**: Distributions of inter-onset interval (IOI) lengths in dances of seven females. IOI numbers per individual are annotated. Species are color-coded: *N. annamensis* – orange; *N. gabriellae* – magenta; *N. leucogenys* – blue; *N. siki* – yellow. **b**: Distribution of rhythmic ratios (*r; n* = 1086) in the total dataset of quantified dances. The red mark indicates the mean rhythmic ratio (= 0.5004). Isochrony is indicated by *r* = 0.5.

### 3.4 RESEARCH SURVEY

We received 29 responses from representatives of 28 institutions to the self-administered survey on crested gibbon dance. Of these, 27 responses came from zoos and rescue centres in Europe, the United States, and Vietnam, and two from researchers studying gibbons in their natural habitats. Respondents, were animal caretakers (48.3%), scientific staff (curators / researchers, 31%), veterinarians (6.9%), and 4 persons who reported to represent neither of the mentioned professions but extensive experience with *Nomascus* gibbons (13.8%). Respondents’ experience with gibbons varied between “several months” and 39 years. The respondents with at least one year of experience with small apes (*n* = 28), had worked with these primates for 10.1 ± 8.8 years on average.

The respondents observed a total of 146 captive gibbons (*N. annamensis* (1 institution), *N. gabriellae* (6 institutions), *N. leucogenys* (20 institutions), *N. siki* (1 institution)) as well as 8 wild gibbon groups (*N. concolor* - 4 family groups, Xiaobahe, Wuliang Mountains, Yunnan, China; *N. nasutus* - 4 family groups, Trung Khan District, Vietnam). Within this extensive sample, 16 respondents (55.2%) have observed dances, which occurred in all covered species except for *N. concolor*, while 13 (44.8%) reported to have never noticed this behaviour. Dancing individuals were exclusively female.

*Nomascus* females of all age classes, except for the youngest (0-3 years) were observed to dance. However, older juveniles and young subadults (3-5 years) were only observed to dance by two respondents (6.9%), with one specifying that dance behaviour onset at an age of 4 years. Dance was still observed in females older than 35 years (4 respondents, 13.8%) and is therefore present in senescent individuals.

Observed dances were not just targeted towards other gibbons (12 responses, 75% of positive responses), but even more so towards humans (13 responses; 81% of positive responses). Three respondents even reported to have only observed human-targeted dances (19% of positive responses). Only one out of all respondents working with captive gibbons communicated that dances were restricted to conspecific communication. Finally, one respondent noted observed dances were also directed towards primates that were neither gibbons nor humans (a situation we observed as well in the female “Lucki” at Zoo Duisburg) and three of the surveyed reported to have seen dances that were not targeted towards a conspecific or heterospecific receiver, possibly representing displacement behaviour.

Gibbon-directed dances were primarily observed in the context of copulation (9 responses, 75% of responses reporting dance targeted at conspecifics) and socialisation / grooming (4 responses, 33%) and to some extent also in stressful situations (2 responses, 17%) or in feeding contexts (1 response, 8%). Human directed dances were most frequently reported from situations involving interspecific socialisation and grooming (9 responses, 69% of responses reporting dance targeted at humans), and contexts of disturbances (5 responses, 38%; possibly displacement actions) as well as feeding (4 responses, 31%). The latter include dances apparently performed in anticipation of receiving food. Analogous to that, one of the respondents reporting dance targeted at humans (8%) reported that dances occurred before commencing a training session. Three respondents (23%) interpreted dances directed towards humans as sexual solicitations. No respondent noticed effects of hormonal contraception on dance behaviour.

Ultimately, three out of all 29 respondents (10.3%) noted that they have observed behaviours similar to crested gibbon dances in female siamangs.

## 4. DISCUSSION

### 4.1 GENERAL DISCUSSION

Our data and accompanying survey demonstrate that dance is a common social display in gibbons of the genus *Nomascus*, also occurring in species so far not reported to exhibit it, and which do not show polygynous mating systems to notable extents (e.g., Barca et al. 2016; Hu et al., 1989; Kenyon et al. 2011; see Fig. 4). In fact, there is now evidence for female dancing in all *Nomascus* species (*N. hainanus*: Li et al. 2022, Zhou et al. 2008; *N. leucogenys*: Lukas et al., 2002, this study; *N. nasutus*: Fan et al., 2016; *N. annamensis*, *N. gabriellae*, *N. siki*: this study), suggesting it to be a shared trait across the genus.

**Fig. 4.**
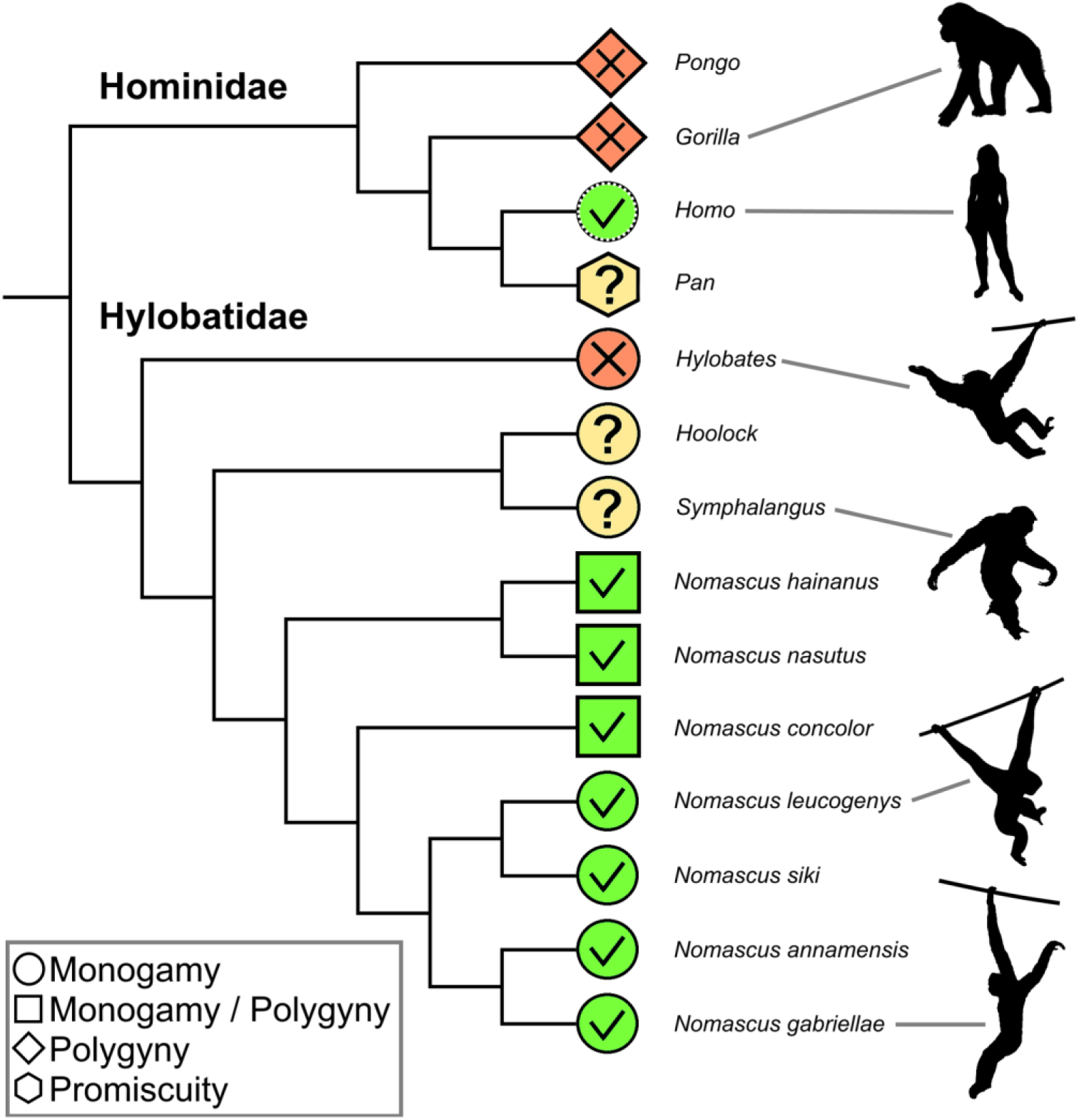
The occurrence of dance in social communication and mating systems across apes (superfamily Hominoidea). Colours and symbols denote the presence (green tick) or absence (red cross) of dancing based on current evidence and the definition applied herein. The genera *Hoolock*, *Pan*, and *Symphalangus* show evidence for rhythmic displays that may potentially qualify as dance but are insufficiently characterised at the moment (yellow question mark). The original human mating system is tentatively hypothesised as a type of monogamy here. Mating systems are defined as: monogamy – both sexes mate primarily with one partner in a given time period; polygyny: males have typically at least two (or multiple) female partners, while females have primarily one male partner; promiscuity – both sexes mate with multiple partners. Hylobatid tree topology follows Shi & Yang, 2018 and Thinh et al. 2010

For the Western black crested gibbon (*N. concolor*), this behaviour has been unequivocally captured by a Chinese TV documentary (CCTV, 2016) whereas available accounts on the mating behaviour of this species, do not mention female displays of any form (Huang et al. 2013; Zheng 1988).

Our data support the notion that only sexually mature female crested gibbons perform dances, but also that dancing is still present in female individuals of advanced age that may no longer be reproductively active. Survey results suggests that the onset of dance behaviour falls within the time window of menarche (4.6 – 7.7 years – Bolechova et al. 2019; Margulis and Hálfdanardóttir 2021) and the subsequent development of an adult pelage colouration in captive *Nomascus* (5 – 8 years; Geissmann et al. 2000), however future studies are required to confirm a correlation of dance onset and these ontogenetic events at the individual level. Furthermore, our findings show that dances in *Nomascus* represent a form of intentional communication, as do visual gestures previously described for great as well as small apes (e.g., Byrne et al. 2017; Liebal et al. 2004). While our survey results confirm that dances act as proceptive signals that solicit copulation (Fan et al. 2016; Li et al. 2022; Lukas et al. 2002; Zhou et al. 2008), they also occur in a number of different behavioural contexts relating to non-sexual arousal, at least in zoo-housed gibbons, and regardless of whether a female is experiencing oestrus or is contracepted (although the frequency of dances in general could be affected by the oestrus cycle – Lukas et al. 2002). Future research needs to clarify whether dancing in certain circumstances is specific to captive gibbons. In orang-utans, for instance, it has previously been shown that gesture use can differ markedly between wild and captive populations (Fröhlich et al. 2021).

Our finding that captive female crested gibbons often direct dances towards human keepers prior to feeding or during social interactions suggests that they can act in non-sexual attention-getting, as originally suggested by Maxwell (1984) and later hypothesised by Fan et al. (2016) (see Caspar et al. 2020 for further discussion on attention-getting in captive gibbons). Alternatively, rather than being goal-directed, dances may simply allow females to relieve tension in context of arousal or frustration (see Botting and Bastian 2019). However, given the intentional dimensions and complex structures of the dances that we described here, the first option appears more plausible. In any case, frustration paired with excitement may be cross-context drivers of dancing. Indeed, stereotyped body-shaking, which was incorporated into 6 of the studied dances (32%), has previously been described as a frustration-related behaviour in gibbons (e.g., Baldwin and Teleki 1976; Maxwell 1984).

We demonstrate that dances follow an isochronous rhythm that this independent of the length of movement intervals and which appears to be conserved across the species studied here. Isochronous communicative behaviours and their evolution are intensely discussed in the primatological community and beyond (Ravignani & Madison, 2017) but so far mostly with a focus on vocalizations (e.g., De Gregorio et al., 2023; Ma et al., 2024) or other acoustic displays (Dufour et al., 2015). Indeed, isochrony has been demonstrated in gibbon songs, including those of *Nomascus* species (De Gregorio et al., 2023; Ma et al., 2024). Apparently, isochronous patterns are not restricted to vocal behaviors in small apes but are also found in their dance displays. To our knowledge, we provide the first confirmation of isochrony in a non-human primate visual display behavior.

Different from any visual gesture described for other non-human primates to date, *Nomascus* dances can display variable grouping structures (another candidate behaviour potentially exhibiting this quality might be the courtship “dance” of gracile capuchin monkeys (*Cebus* sp. - see e.g., Perry, 1996) which remains understudied). Two or more structural groups could be identified in 71% of the dance displays analysed. It remains to be seen if, and how grouped dances differ from homogeneous dances with regards to their meaning. Notably, in addition to one-level grouping structures, we identified candidates for two-level grouping structures in the shape of nested groups, as proposed in linguistic approaches to other cognitively complex behaviours. Previous work has reported hierarchical structures in the vocal communication of birds (Berwick et al. 2011) and orang-utans (Lameira et al. 2024). Based on the sample size, data collection methods and contexts, it remains unclear whether gibbon dances consist of compositional semantic contents, or iconic components. Interestingly, other complex visual displays exist in non-primate animals, which may also bear a potential hierarchical structure. This is for instance the case of the courtship displays of some birds-of-paradise species of the genus *Astrapia* (Aves: Paradisaeidae) in which one movement pattern, the “flick-pivot” motif, involves repeated wing flicks displayed while the male moves side-to-side in space (Scholes et al. 2017).

### 4.2 PHYLOGENETIC PERSPECTIVES

Because dances, at least in wild crested gibbons, most obviously function in soliciting copulation (Fan et al. 2016; Li et al. 2022; Zhou et al. 2008), they might have evolved as a proceptive gesture. Hence, proceptive displays in other gibbons might provide clues about the evolution of dance behaviour in gibbons.

Nothing resembling a dance is known from dwarf gibbon social communication (genus *Hylobates*; see e.g., Baldwin and Teleki 1976; Palombit 1992; but note dance-like decoy displays in wild Kloss gibbons (*Hylobates klossii*) – Dooley and Judge 2015). For wild Eastern hoolock gibbons (*Hoolock leuconedys*) “head nodding” and branch shaking, have been described as female gestures to solicit mating (Kumar and Sharma 2017) but available literature reports certainly do not suggest that these signals qualify as dance. Nevertheless, via our survey, we received a short video of a female Eastern hoolock housed at the Gibbon Conservation Centre in California, which engages in bobbing movements of the upper body, while socializing with her male partner. This type of display is likely not idiosyncratic, since it has also been observed in other adult females at the Gibbon Conservation Centre (G. Skollar, pers. comm.), which is one of the few places where captive hoolock gibbons are housed. Further and more systematic observations are needed to sufficiently characterise the occurrence of this behaviour in hoolocks. More robust evidence for rhythmic socio-sexual displays is available for siamangs (genus *Symphalangus*). Adult females of this species may engage in jerking movements of the upper body mediated by repeated angling and stretching of the arms, which have been observed both in the wild (“upward-thrust” – Palombit 1992) and in captivity (“jerking body movements” – Liebal et al. 2004; Orgeldinger 1999). This display appears to be very similar to homogeneous or simple dance displays in *Nomascus* and three respondents to our survey actually reported having observed “dances” in siamangs. In adult siamangs, jerking movements have not been reported outside of sexual contexts, so far, but different from *Nomascus*, juvenile siamangs may use them to initiate play (Liebal et al. 2004). Unfortunately, further comparisons between such displays in crested gibbons and siamangs are unfeasible at the moment, because they have never been studied systematically in the latter.

Future research needs to clarifiy if “jerking body movements” of some form represent homologous social signals in the small ape genera *Nomascus*, *Symphalangus,* and potentially *Hoolock*. Interestingly, gibbon phylogenies based on genomic datasets suggest a derived clade formed by these three genera, with *Hylobates* branching off earlier (Carbone et al. 2014; Shi and Yang 2018; Fig. 4). However, given the notoriously conflicting evidence on the topology of the hylobatid family tree (Roos 2016), future studies need to consolidate this phylogenetic hypothesis.

An important question with regards to dance behaviour in gibbons is whether it could be phylogenetically linked to the origins of dance and dance-like gestures in humans (Fan et al., 2016). In light of the current evidence, we see no compelling evidence for this idea. First, the phylogenetic distance between humans and hylobatids in combination with the scarcity or absence of reports on dance behaviour in the non-human great apes and basal-branching gibbons of the genus *Hylobates* argues against a phylogenetic continuity. Second, the uniform structure of gibbon dances across species together with the fact that they appear to be tied to female sexual maturity suggests them to be importantly determined by innate factors, different from human ones. Interestingly, whereas dance in humans is almost exclusively a social endeavour rooted in imitation and the entrainment of movements within a group (e.g., Laland et al., 2016), no indication of this is evident in gibbons. We suggest that human and hylobatid dance, although perhaps based on shared perceptive and sensorimotor principles (see below), originated independently from one another.

### 4.3 FUTURE PERSPECTIVES

This research and the small number of other available studies on dances in crested gibbons are an initial starting point when it comes to elucidating this behaviour. Next steps should address the investigations of the variability of dances at the individual as well as species level, and studies testing for correlates of their structure and frequency. This includes the question whether the different dance types described herein are used to communicate distinct semantic information or signal intensities. Fan et al. (2016) suggested that dances could be a sexually selected behaviour. If so, we would expect that certain aspects of a dance reflect a female’s reproductive fitness. In this context, it is important to address why some females dance, while a substantial percentage appears not to, and what causes individuals to stop dancing at some point in their lives. For instance, de Vries (2004) documented a high frequency of dances (70 instances in 28h of observation) in the female Kanak at Apenheul Primate Park in Apeldoorn (*N. leucogenys*; born 1993; ZIMS GAN: MIG12-29829865), which has not been observed to dance in the last 15 years according to our survey response from the corresponding institution.

Other relevant questions concern the structural analysis of dance movements in humans (see, in particular, Charnavel, 2019) which has focused on perception, specifically, the idea that humans’ perception of dance is shaped by gestalt principles (Wertheimer 1938, and subsequent work). The underlying intuition is that human perception organises a dance sequence into *groups* based on similarity of the movements within a group; significant changes in the movement patterns (such as a change of bodily orientation) will then give rise to the perception of group boundaries (labelled GPRs by Charnavel, 2019, following Lerdahl & Jackendoff’s, 1983, work on music). As of now, we have a very limited understanding of gibbon cognition (see, e.g., King 2021 for recent discussion), which leaves it unclear whether gibbons exhibit gestalt perception as well (see, e.g., Hopkins and Washburn, 2002 on gestalt perception in chimpanzees vs. its absence in rhesus monkeys) and thus how they perceive the dances of their conspecifics.

## Supplementary material

**Supplementary File 1**: Exemplary videos of a complex (time stamp 00:00:00.000), simple (time stamp 00:01:37.500) and homogeneous dance (time stamp 00:02:29.567), combined into a single video.

**Supplementary File 2**: Dataset used in the rhythm analysis, including inter-onset intervals (IOI) and rhythmic ratios (r).

**Supplementary File 3**: List of questions included in the survey

**Supplementary File 4**: Detailed time-stamped description of a dance by a Southern white-cheeked gibbon female (Doremon; EPRC), corresponding to the complex dance display shown in Suppl. File 1.

## Funding

This research was funded by the Career development programme, University of Oslo, EU Horizon 2020 Marie Skłodowska-Curie R&I program, under grant agreement no 945408, and RFIEA+ LABEX, French national grant, ANR-11-LABX-0027-01 (PI Patel-Grosz) and by the European Research Council (ERC) under the European Union’s Horizon 2020 research and innovation program (grant agreement No 788077, Orisem, PI: Schlenker). This research was conducted in part at DEC, Ecole Normale Supérieure - PSL Research University. DEC is supported by grant FrontCog ANR-17-EURE-0017.

## Credit Statement

Conceptualisation: PPG; Biological methodology: KRC, CC; Linguistic methodology: PPG, CC; Data coding: CC, PPG, KRC; Formal analysis: PPG, CC, KRC; Survey: KRC; Resources: KRC, PPG; Data Curation: KRC; Figures: KRC, PPG, CC; Writing – Original Draft: CC, KRC; Writing – Review & Editing: CC, KRC, PPG; Funding: CC, PP.

## Data availability

All videos for which we have permission to share can be found in the following Open Science Framework repository: https://osf.io/cv3a9/

## Acknowledgements

We are indebted to the primate researchers and keepers who send in video material of dance displays in gibbons. First and foremost, we are extremely grateful to Kaylen Kilfeather for numerous high-quality videos from the EPRC which were essential for conducting this study. Further videos were very kindly provided by Nene Haggar, Alexandre Petry, Joanna Husby, Fabian Ohl, Katja Rudolph, Julia Ruiz Laguna, Elke Schwierz, Gabriella Skollar, and Saskia Winking, some of which will only be used in future studies.

We thank the Gibbon Taxon advisory group (TAG) of the European Association of Zoos and Aquaria (EAZA) and the Gibbon Species Survival Plan® (SSP) steering committee of the Association of Zoos and Aquariums (AZA) for their support of the study and especially Hélène Birot, Susan Margulis, Gina Munir, and Holly Thompson for facilitating communication and data collection.

We wish to acknowledge all institutions that shared their experiences with gibbon dance through the survey: Apenheul Primate Park (Apeldoorn, Netherlands), Bioparc Doue la Fontaine, Centro de Conservación Zoo Córdoba, CERZA Zoo (Lisieux, France), Chattanooga Zoo, Cheyenne Mountain Zoo (Colorado Springs, Colorado), Disney’s Animal Kingdom (Bay Lake, Florida), Durrell Wildlife Conservation Trust (Jersey, UK), Edinburgh Zoo, Endangered Primate Rescue Centre, Gibbon Conservation Center, Gdański Ogród Zoologiczny (Gdansk, Poland), Liberec Zoo, Lyon Zoo, Ostrava Zoo, Paradise Wildlife Park (Broxbourne, UK), Parc animalier de Sainte-Croix (Rhodes, France), Parc de Branféré, Perth Zoo, Parken Zoo (Eskilstuna, Sweden), Pittsburgh Zoo, Virginia Zoo, Wildlife Sarai (Winston, Oregon), Zoo Łódź, Zoo Planckendael (Mechelen, Belgium), Zoo Ústí nad Labem.

We are grateful to Sabine Begall, Marléne Baumeister and Fabian Pallasdies for advice on statistics and Heike Kessels for data coding to assess inter-rater reliability.

Finally, we thank Le-Jia Zhang for translating Chinese references and for making us aware of a CCTV documentary containing footage of dance displays in *Nomascus concolor*.

## Notes

https://osf.io/cv3a9/

### Competing Interest Statement

The authors have declared no competing interest.

