## Supplementary File 3 for "Dance displays in gibbons: Biological and linguistic perspectives on structured, intentional and rhythmic body movement"

**1) What is your associated institution / field site? (optional: You may want to also provide your name)**

Free-form text response

**2) What is your position at your institution?**

<u>Options:</u>

Researcher / Curator

Veterinarian

Animal caretaker

other (free-form text response)

**3) How long did you monitor / work with gibbons at your institution / site?**

Free-form text response

**4) How many gibbons did you opportunistically or systematically monitor? Please provide the species and sex of the respective individuals, or give approximations.**

Free-form text response

**5) In how many individuals did you observe "dances"? Please also note the sex of the respective individuals.**

Free-form text response

**6) Which was the (approximate) age range of individuals that were observed to "dance"?**

<u>Options:</u>

0 - 3 years

3 - 5 years

5 - 7 years

7 - 10 years

10 - 25 years

25 - 35 years

35 + years

Not applicable

**7) Have you observed "dances" directed towards ...**

<u>Options:</u>

other gibbons

humans

other primate species (neither gibbons nor humans)

self (displacement behavior; no receiver of the dance was apparent)

I have never observed "dances"

**8) In which context have you observed "dances" directed towards a gibbon?**

Options:

invitation to copulate

socializing / grooming

stressful situation / disturbances

feeding / foraging

other (free-form text response)

**9) In which context have you observed "dances" directed towards a human?**

Options:

invitation to copulate

socializing / grooming

stressful situation / disturbances

feeding / foraging

other (free-form text response)

**10) Did you notice effects of hormonal contraception on the occurence of "dances" in captive female gibbons? If so, please provide details.**

Free-form text response

**11) Are there any additional observations on gibbon "dance" behavior that you want to share (e.g. frequency of occurence, uniformity of dance patterns)?**

Free-form text response

**12) Do you have video material of "dances" that you would be willing to share with us for further study (the videos do not need to be recorded at the institution/ field site that you are currently associated with) and / or are you interested in the results of our study? If so, please provide your name and an email address so we can contact you.**

Free-form text response
