## Supplementary File 4 for "Dance displays in gibbons: Biological and linguistic perspectives on structured, intentional and rhythmic body movement"

Table S1 provides a full analysis of the complex dance that underlies Fig. 1c, which corresponds to the complex dance in Supplementary File 1. ELAN 6.1 was used to establish time stamps.

Supplementary File 1 includes three dances; the complex dance is preceded by a black title screen shown for a time span of 00:03.799, which is not included in the time stamps below; i.e., time stamp 00:03.981 in Table S1 corresponds to 00:08.780 in Supplementary File 1. A version of the video without the black title screen, where the time stamps match the ones given in Table S1 can be downloaded from https://osf.io/u4ng6

For expository simplicity, Fig. 1c is a schematic representation of the dance up to 01:09. Note that the distance between twitches increases in the part from 01:15.782 until 01:30.432, indicating that this may not be part of the dance proper, e.g., due to the gibbon being disrupted or distracted.

Table S1: Behavioural variables annotated in the complex dance illustrated in Fig. 1c

| Grouping by direction | Grouping by posture | Time stamp annotation |
|---|---|---|
| Left-Right | Sitting (Crouching) | 00:03.981 - assume **start** position (posture: crouching) |
| | | 00:05.048 - onset of movement towards the **right** (posture: crouching) |
| | | 00:06.008 - onset of movement towards the **left** (posture: crouching) |
| | | 00:07.408 - onset of movement towards the **right** (initial posture: crouching; end posture: sitting) |
| | | 00:08.269 - onset of movement towards the **left** (posture: sitting) |
| Up-Down | N/A | 00:09.329 - onset of **upwards** movement (from sitting to standing) |
| | | 00:10.209 - onset of **downward**s movement (from standing to sitting) |
| | | 00:11.009 - onset of **upwards** movement (from sitting to standing) |
| | | 00:11.679 - onset of **downwards** movement (from standing to sitting) |
| | | 00:12.679 - onset of **upwards** movement (from sitting to standing) |
| | | 00:13.829 - onset of **downwards** movement (from standing to sitting) |
| | | 00:15.274 - onset of movement towards the **left** (posture: sitting) |
| | | 00:16.011 - onset of **upwards** movement (from sitting to standing) |
| | | 00:17.156 - onset of **downwards** movement (from standing to sitting) |
| | | 00:18.116 - onset of **upwards** movement (from sitting to standing) |
| | | 00:18.936 - onset of **downwards** movement (from standing to sitting) |
| | | 00:19.966 - move towards the **right** into a resting position for a pause (posture: sitting) |
| | | 00:23.216 - return to **upright** position, continue pause (posture: sitting) |
| | | 00:24.156 - onset of short **up-down** movement (posture: sitting to crouching to sitting) |
| | | 00:25.356 - onset of **upwards** movement (from sitting to standing, leaning leftward) |
| | | 00:26.626 - onset of **downwards** movement (from standing to crouching, leaning leftward) |
| Left-Right | Sitting | 00:27.386 - onset of movement towards the **right** (initial posture: crouching; end posture: sitting) |
| | | 00:28.316 - onset of movement towards the **left** (posture: sitting) |

| | | |
|---|---|---|
| | | 00:29.116 - onset of movement towards the **right** (posture: sitting) |
| | | 00:29.996 - onset of movement towards the **left** (posture: sitting) |
| | | 00:30.896 - onset of movement towards the **right** (posture: sitting) |
| | | 00:31.819 - onset of further movement more towards the **right** (posture: sitting) |
| | | 00:32.859 - onset of **upwards** movement (from sitting to standing, towards the left) |
| | Standing | 00:34.009 - onset of movement towards the **right** (posture: standing) |
| | | 00:34.909 - onset of movement towards the **left** (posture: standing) |
| | | 00:35.709 - onset of movement towards the **right** (posture: standing) |
| | | 00:36.599 - onset of movement towards the **left** (posture: standing) |
| | | 00:37.399 - onset of movement towards the **right** (posture: standing) |
| | | 00:38.199 - onset of movement towards the **left** (posture: standing) |
| | | 00:38.859 - onset of movement towards the **right** (posture: standing) |
| | | 00:39.569 - onset of movement towards the **left** (posture: standing) |
| | | 00:40.519 - onset of movement towards the **right** (posture: standing) |
| | | 00:41.279 - onset of movement towards the **left** (posture: standing) |
| | | 00:42.169 - onset of movement towards the **right** (posture: standing) |
| | | 00:42.959 - onset of movement towards the **left** (posture: standing) |
| | | 00:43.809 - onset of movement towards the **right** (posture: standing) |
| | | 00:44.539 - onset of **downwards** movement (from standing to sitting, leaning rightward) |
| | Sitting | 00:45.369 - onset of movement towards the **left** (posture: sitting) |
| | | 00:47.149 - onset of movement towards the **right** (posture: sitting) |
| | | 00:48.079 - onset of movement towards the **left** (posture: sitting) |
| | | 00:48.889 - onset of further movement more towards the **left** (posture: sitting) |
| | | 00:50.609 - onset of movement towards the **right** (posture: sitting) |
| | | 00:51.539 - onset of movement towards the **left** (posture: sitting) |
| | | 00:52.179 - onset of movement towards the **right** (posture: sitting) |
| | | 00:53.049 - onset of movement towards the **left** (posture: sitting) |
| | | 00:53.819 - onset of further movement more towards the **left** (posture: sitting) |
| | | 00:54.819 - onset of movement towards the **right** (posture: sitting) |
| | | 00:55.849 - onset of movement towards the **left** (posture: sitting) |
| | | 00:56.719 - onset of **upwards** movement (from sitting to standing) |
| | Standing | 00:57.689 - onset of movement towards the **right** (posture: standing) |
| | | 00:58.419 - onset of movement towards the **left** (posture: standing) |
| | | 00:59.159 - onset of movement towards the **right** (posture: standing) |
| | | 00:59.919 - onset of movement towards the **left** (posture: standing) |
| | | 01:00.659 - onset of movement towards the **right** (posture: standing) |
| | | 01:01.559 - onset of movement towards the **left** (posture: standing) |
| | | 01:02.449 - onset of further movement more towards the **left** (posture: standing) |
| | | 01:03.099 - onset of movement towards the **right** (posture: standing) |
| | | 01:03.897 - onset of further movement more towards the **right** (posture: standing) |
| | | 01:04.952 - onset of movement towards the **right** (initial posture: standing; end posture: crouching) |
| | | 01:05.662 - onset of movement towards the **left** (initial posture: crouching; end posture: standing) |
| | | 01:06.552 - onset of movement towards the **right** (initial posture: standing; end posture: crouching) |
| | | 01:07.302 - onset of movement towards the **left** (initial posture: crouching; end posture: standing) |
| | | 01:08.262 - onset of movement towards the **right** (initial posture: standing; end posture: crouching) |

| | | |
|---|---|---|
| | | 01:09.092 - onset of movement towards the **left** (posture: crouching) |
| Up-Down | N/A | 01:09.702 - onset of **upwards** movement (from sitting to standing) |
| | | 01:12.822 - onset of **downwards** movement (from standing to sitting) |
| | | 01:13.732 - onset of **upwards** movement (from sitting to standing) |
| | | 01:14.872 - onset of **downwards** movement (from standing to sitting) |
| (Left-Right) | Sitting | 01:15.782 - onset of movement towards the **left** (posture: sitting), followed by more relaxed movement and pause |
| | | 01:19.192 - onset of movement towards the **right** (posture: sitting) |
| | | 01:22.332 - onset of **upwards** movement (from sitting to standing, leaning rightward) |
| | Standing (Crouching) | 01:23.162 - onset of movement towards the **right** (initial posture: standing; end posture: crouching) |
| | | 01:25.312 - onset of movement towards the **left** (initial posture: crouching; end posture: standing) |
| | | 01:26.292 - onset of further movement more towards the **left** (posture: standing) |
| | | 01:27.222 - onset of further movement more towards the **left** (initial posture: standing; end posture: crouching) |
| | | 01:28.142 - onset of movement towards the **right** (initial posture: crouching; end posture: standing) |
| | | 01:29.282 - onset of further movement more towards the **right**, lifting one leg (posture: standing) |
| | | 01:30.432 - **end** of dance, relaxation of body posture and move away |
